# PRPF31 interacts with PRPH2 confirmed by co-immunoprecipitation and co-localization

**DOI:** 10.1101/2022.06.26.497680

**Authors:** Xiaoqiang Xiao, Fangyi Ling, Chongbo Chen, Jiajian Liang, Yingjie Cao, Yanxuan Xu, Haoyu Chen

**Author notes:** Co-responding author: Haoyu Chen. These authors contribute equally to the manuscript (Xiaoqiang Xiao, Fangyi Ling).

## Abstract

**Purpose:** To investigate the potential interaction between PRPF31 and PRPH2.

**Methods:** HEK293T and human retinal pigment epithelial cells 19 (APRE-19) were used for our experiments. eGFP and mCherry reporter expression vectors were constructed for PRPF31 and PRPH2, respectively. Immunoblotting and co-immunoprecipitation (Co-IP) were used for gene expression validation and protein interaction. Immunofluorescence staining assay was used to test the co-localization analysis of PRPF31 and PRPH2.

**Results:** PRPF31-eGFP and PRPH2-mcherry were highly expressed in HEK293T and APRE-19 cells on fluorescence microscopy and western blot. Co-IP experiments showed that PRPF31 could be pulled down with an anti-PRPH2 antibody. There was co-localization between PRPF31 and PRPH2 in HEK293T, APRE-19 and mouse retina.

**Conclusion:** Co-IP and co-localization experiments suggest that PRPF31 interacted with PRPH2.

## Introduction

Retinitis pigmentosa (RP) is a group of inherited disorders characterized by photoreceptor and retinal pigment epithelium dystrophy.[1] The patients with RP usually complain of trouble seeing at night and decreasing peripheral vision. Central visual acuity may also be involved in end-stage or severe types.[2] With the development of genetic techniques, 69 genes have been identified as the causative gene for RP.[3-6] Rhodopsin is the earliest identified gene for RP.[7, 8] The rhodopsin gene encodes a principal protein of rod photoreceptor outer segments, which is the critical protein in phototransduction. The light is absorbed by rhodopsin and causes isomerization of 11-cis-retinal to all-trans-retinal. And all-trans-retinal is converted back into 11-cis retinal before re-entry into the cycle.[8] The dysfunction of rhodopsin and other proteins associated with the visual cycle will cause retinal dystrophy and visual impairment.

Peripherin2/rds (PRPH2) is expressed in the rim regions of the flattened disks that contain rhodopsin in the outer segment of the photoreceptor. It functions as an adhesion molecule involved in the stabilization and compaction of outer segment disks.[9] PRPH2 was reported to couple rhodopsin to the CNG channel in the outer segments of the rod.[10] Pre-mRNA processing factor 31 homolog, PRPF31, is widely expressed in many tissues of the body. However, the mutation of PRPF31 only causes RP.[11] It was reported that PRPF31 could bind to small nuclear ribonucleoprotein, snRNP, U5 and U4/U6 and regulate the processing of rhodopsin pre-mRNA.[12]

Both PRPF31 and PRPH2 are the causative genes for retinitis pigmentosa. And both of them are associated with the balance of rhodopsin. In this study, we aim to investigate the co-expression and interaction of PRPF31 and PRPH2.

## Methods

### Cell culture

HEK293T cells and ARPE-19 cells were cultured in Dulbecco’s modified Eagle’s medium (DMEM) supplemented with 10% FBS (BI, Cat.04-001-1ACS) and 1% Penicillin / Streptomycin (Gibco, Cat 15140122) at 37 °C with 5% CO2.

### Plasmids construction

PRPF31-eGFP vectors were constructed by Genomeditec (Shanghai, China). Lentivirus was generated from Gene Pharma (Shanghai, China). Briefly, PRPH2 expression plasmid was constructed by inserting the ORF (open reading frame) sequence into a lentivirus-specific vector and fused to a mCherry fluorescence gene, then packed into lentivirus (Gene Pharma, Shanghai, China). PRPF31 ORF sequence, as well as 3Flag tag sequence, was inserted into pcDNA3.1-CMV containing a neomycin resistance gene (Gene Pharma).

### Transient transfection

HEK293T and ARPE-19 cells were transfected with Lipofectamine 2000 reagent (Invitrogen) according to the manufacturer’s instructions. Transfection conditions were maintained for 6 hours, and the cells were washed twice with PBS and returned to the full culture medium. Transgene-expressing cells were confirmed by immunoblotting.

### Immunoblotting

HEK293T and ARPE-19 cells were washed once with cold PBS, and subsequently lysed in 1xRIPA (50 mM Tris/pH 7.4, 150 mM NaCl, 1% Triton X-100, 1% sodium deoxycholate, 0.1% SDS, sodium orthovanadate, sodium fluoride, EDTA, leupeptin, and protease inhibitors) via three cycles of liquid nitrogen /37°C water bath. Lysates were centrifuged at 15,000g for 15 min at 4 °C and quantified by the Bradford Protein Assay (Thermo Scientific). SDS-PAGE gel was used for protein electrophoresis. Five percent milk prepared with 1×TBST was used for membrane block at room temperature for 1 hour. Primary antibodies, including anti-Flag (1:1000, Sigma), and anti-PRPH2 (1:500, Abcam), were incubated with the WB membrane at 4&C overnight. Then, the membranes were washed with TBST and incubated with peroxidase-conjugated secondary antibodies (1:5000, Cell Signaling) for 1 hour and detected with enhanced chemiluminescence (Tanon).

### Immunofluorescence Staining

For co-localization analysis, HEK293T cells were cultured and co-transfected with PRPF31-eGFP and PRPH2-mCherry vectors for 24 hours. Then cells were fixed and permeabilized, washed, and counter-stained with DAPI. Cells were analyzed with a confocal microscope (SP5, Leica Biosystems).

### Co-Immunoprecipitation (CO-IP)

HEK293T and APRE-19 cells were cultured in a 100mm-culture dish. Vectors PRPF31-FLAG and PRPH2-Cherry were co-transfected into these cells with lipofectamine 2000. After 30 hours of transfection, cells were washed with cold PBS, followed by lysed using 1×NP40 lysis buffer supplemented with protease inhibitors Coaktails (Roche) and phosphatase inhibitors(Roche). After centrifuged at 15,000×g for 15 min at 4 °C, per 100 μL of the supernatant was added with 10 μL Protein A/G PLUS-Agarose (Santa Cruz, CA, USA) and incubated with the Anti-Flag antibody overnight at 4 °C. The next day, the immunoprecipitants were washed five times and boiled with 2×SDS Loading Buffer. Do western blotting assay according to the protocol above. For co-IP assay in APRE-19, APRE-19 were cultured in a 100 mm dish for 24 hours and then lysed with 1×NP40 lysis buffer supplemented with protease inhibitors Coaktails (Roche) and phosphatase inhibitors(Roche). The supernatants were added IgG and Protein A/G beads for pre-clearance after centrifugation(15000g,15min,4&C). Then, Anti-PRPH2 antibody and protein A/G beads were added to the pre-cleared supernatant and incubated overnight at 4 °C. Collected the protein A/G beads by centrifugation after 5 times wash with lysis buffer and used for the immunoblotting assay.

## Results

Reporter gene assay with fluorescence microscopy showed that PRPF31-eGFP and PRPH2-mCherry were successfully transfected into the HEK293T cells (Figure 1A-D). The strong fluorescence signals from eGFP (green) and mCherry (Red) confirmed that the expression vectors for PRPF31 and PRPH2 highly expressed the target proteins. Further study validation of the expression was confirmed by immunoblotting assay (Figure 2E).

**Figure 1.**
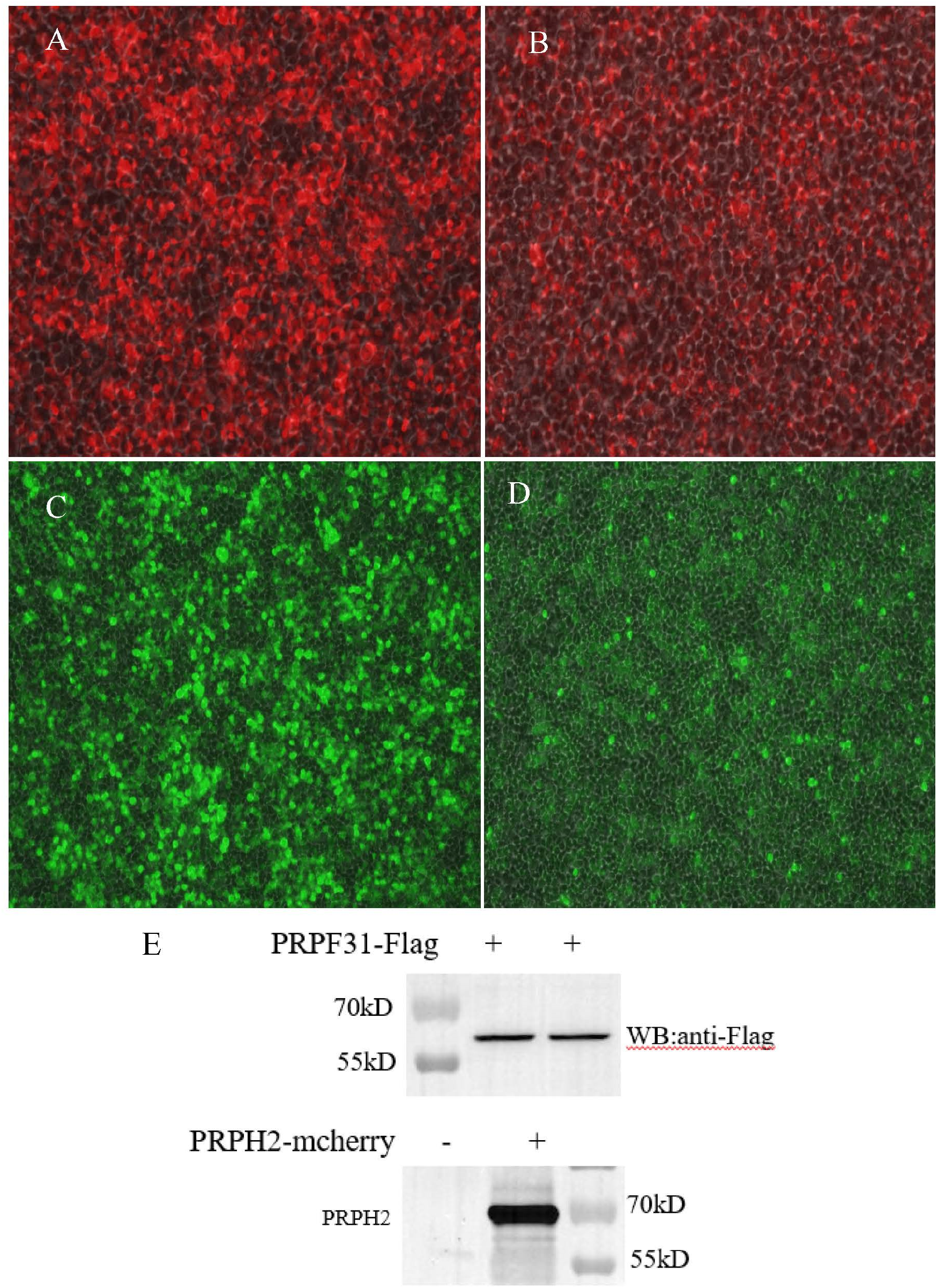
Plasmids of PRPF31 and PRPH2 were successfully transfected into HEK293T cells. A. PRPH2-mcherry. B Control-mcherry. C. PRPF31-eGFP. D. Control-eGFP. E. Immunoblotting showed both PRPF31-Flag and PRPH2-mcherry were detected.

**Figure 2.**
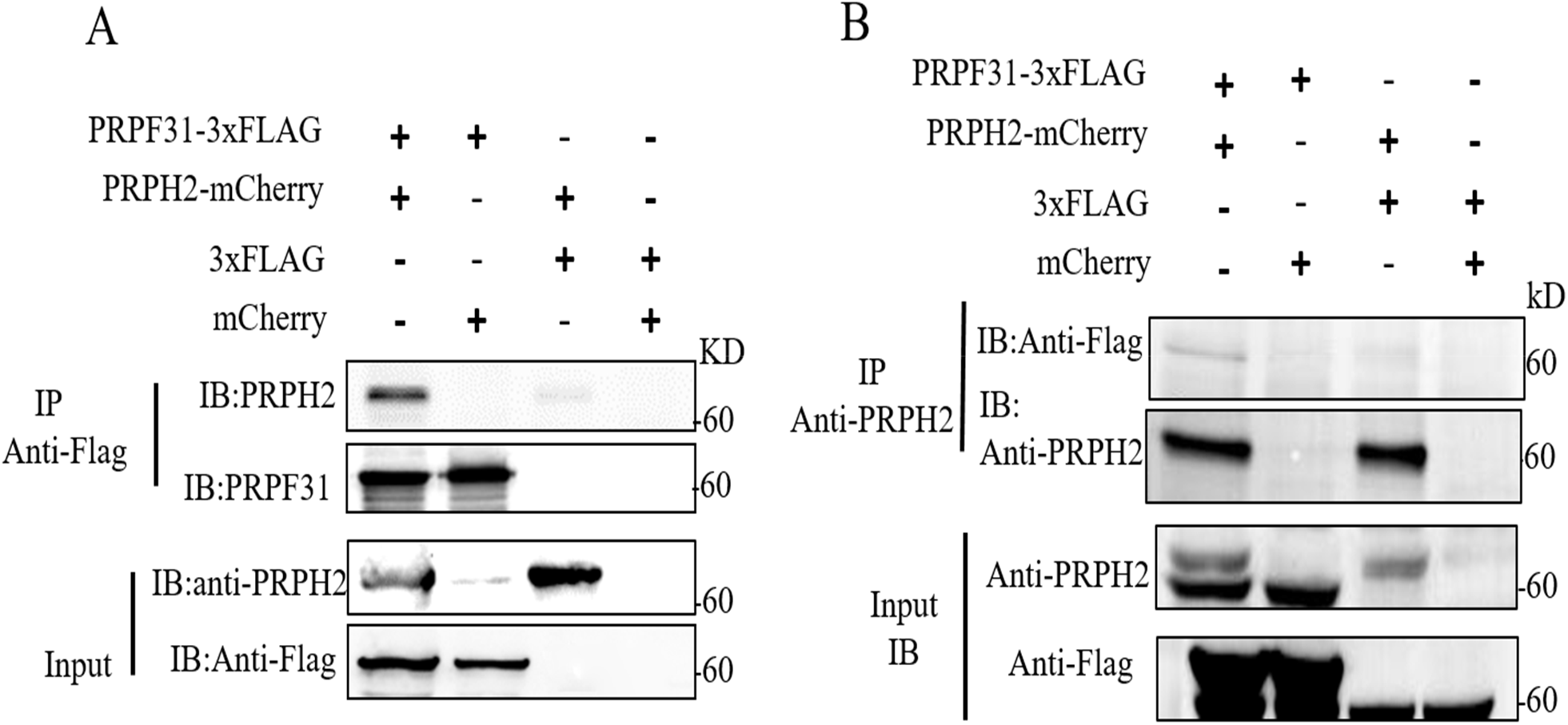
PRPF31 interacts with PRPH2 in HEK293T cell via Co-immunoprecipitation (Co-IP) assay. CoIP was performed using anti-Flag antibody (a) or anti-PRPH2 antibody (b). The upper panel showed IP results and the lower panel showed Input results. IB, immunoblotting, 3×Flag, Empty vector for PRPF31 expression vector, mCherry, empty vector for PRPH2-mCheryy expression vector.

To test the interaction between PRPF31 and PRPH2, we co-transfected the two vectors PRPF31-3Flag and PRPH2-mCherry into HEK293T cells. Co-IP and immunoblotting assay showed that PRPH2 could be pulled down using the anti-Flag antibody (Figure 2A). Moreover, using an anti-PRPH2 antibody as bait for Co-IP assay, we also observed a weak immunoblotting PRPF31 band in HEK293T co-expressed PRPF31-3Flag and PRPH2-mCherry (Figure.2B) In ARPE-19 cells, the co-IP results showed that PRPF31 could be pulled down using the anti-PRPH2 antibody(Figure.3).

**Figure 3.**
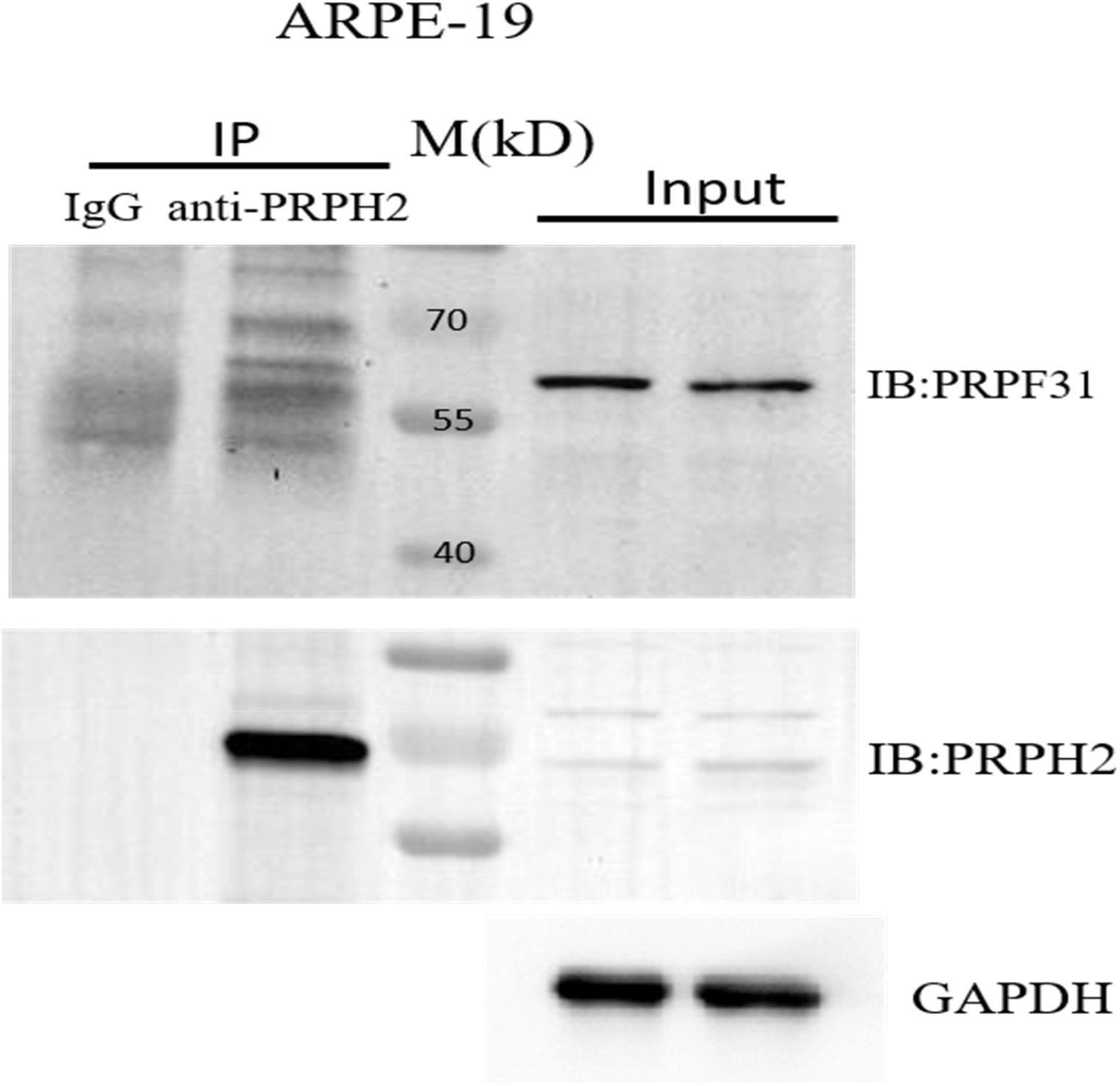
PRPF31 interacts with PRPH2 in ARPE-19 cells via Co-Immunoprecipitation (IP) assay. CoIP was performed using an anti-PRPH2 antibody. IgG was used as a negative control. GAPDH is also used to check the total proteins in IP. The upper panel is the IP results pull-downed by anti-PRPH2 antibody, and the same member was used for Immunoblotting (IB) for the PRPH2 test (the lower panel).

The results of laser confocal microscopy showed that PRPF31-eGFP and PRPH2-m Cherry partially co-localized in the cytoplasmic area of HEK293T cells (Figure 4) and APRE-19 (Figure 5). Immunohistochemistry showed that PRPF31 and PRPH2 partially co-localized in the photoreceptor of C57BL/6J mice (Figure 6).

**Figure 4.**
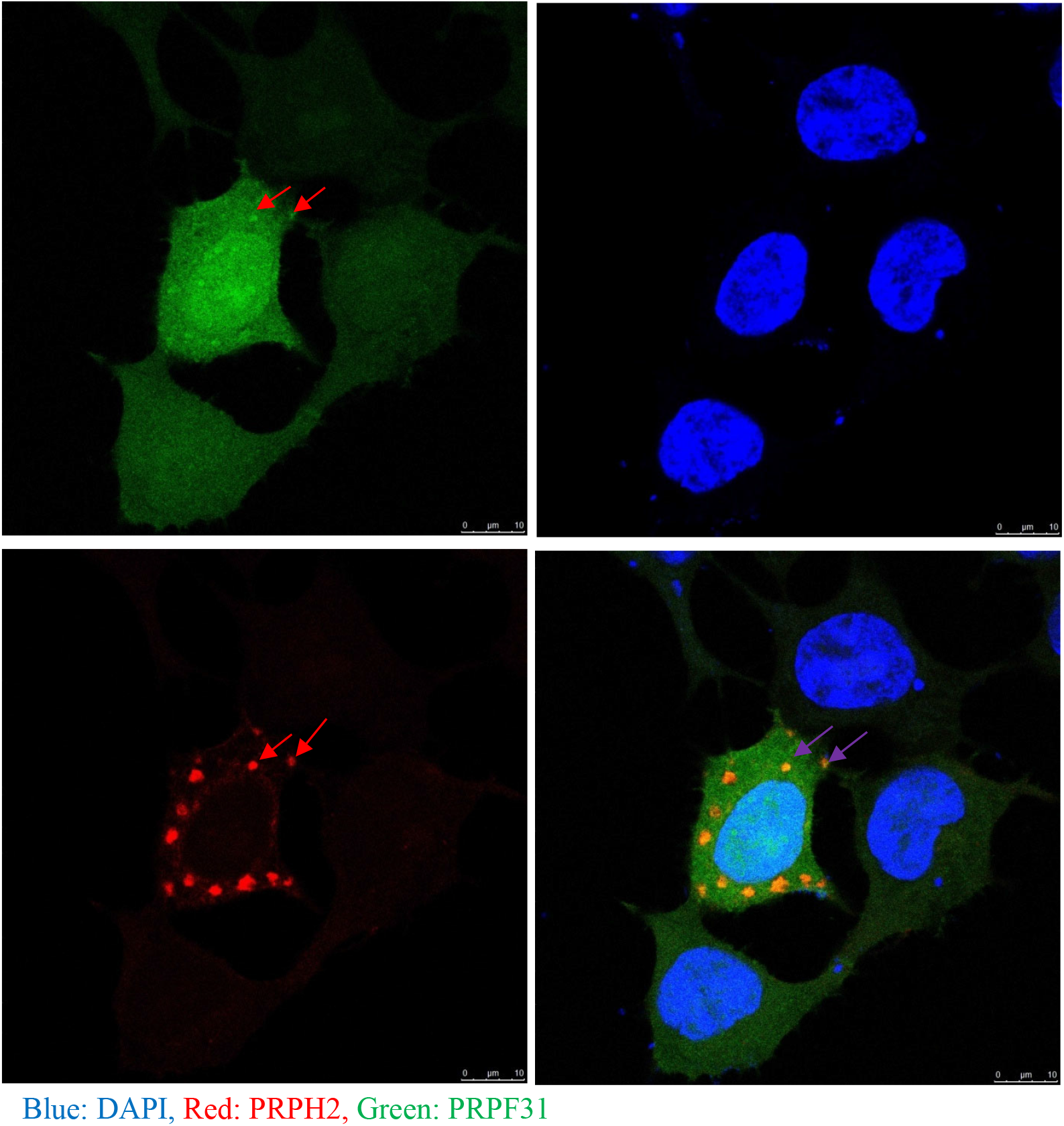
PRPF31 and PRPH2 can partially co-localize in HEK293T cells. PRPF31-eGFP and PRPH2-Cherry were co-transfected into HEK293T cells. Images were obtained at 24h post-transfection with a laser confocal microscope. Red arrows showed the co-localization sites. DAPI was used to stain the nucleus

**Figure 5.**
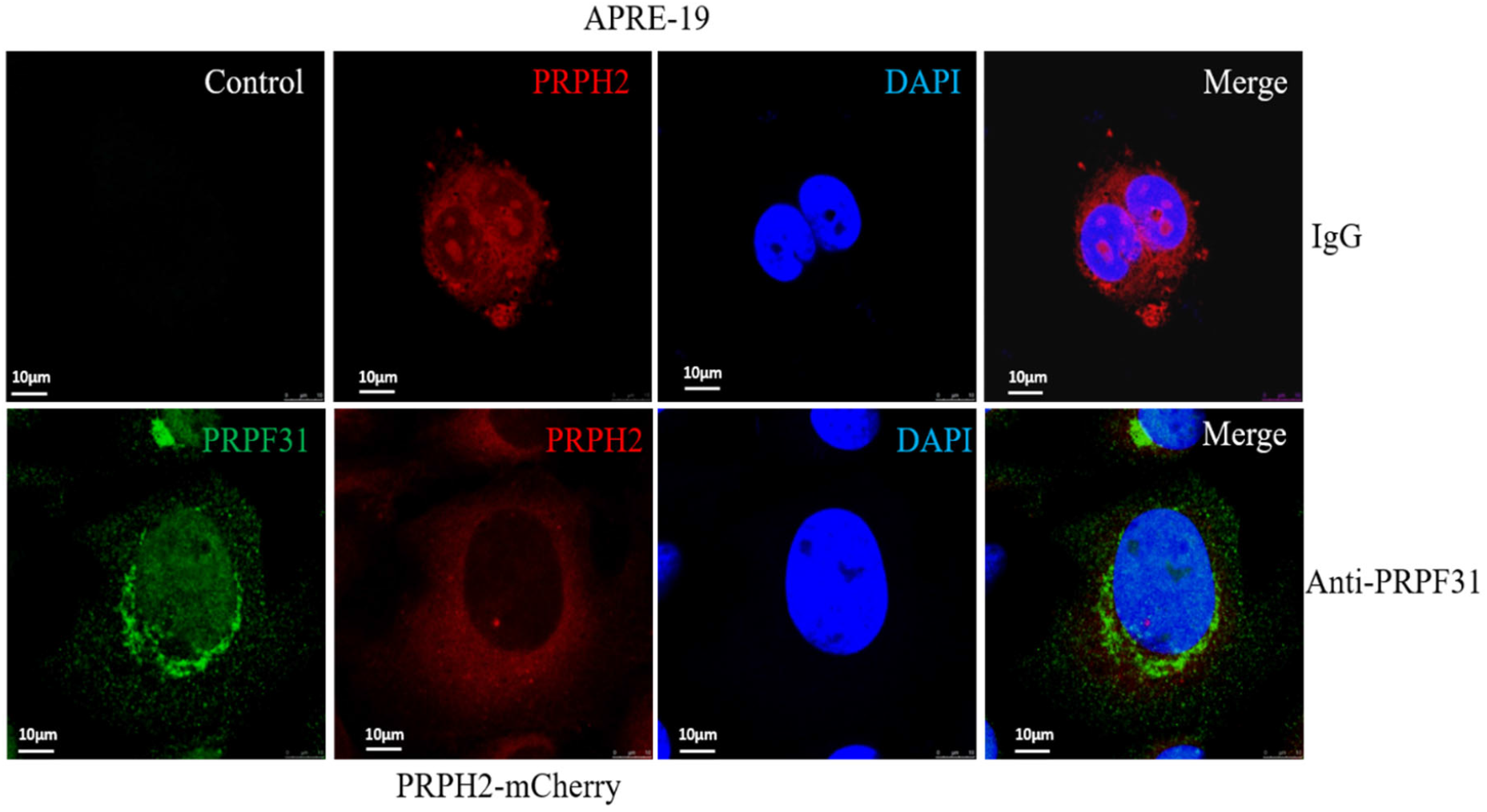
PRPF31 and PRPH2 can partially co-localize in the ARPE-19 cell. PRPH2-Cherry was transfected into ARPE-19 (red). IgG (the upper panel) and anti-PRPF31 antibody were used for immunocytochemistry (green) (The lower panel). DAPI was used to stain the nucleus (blue). Images were taken via laser confocal microscope.

**Figure 6.**
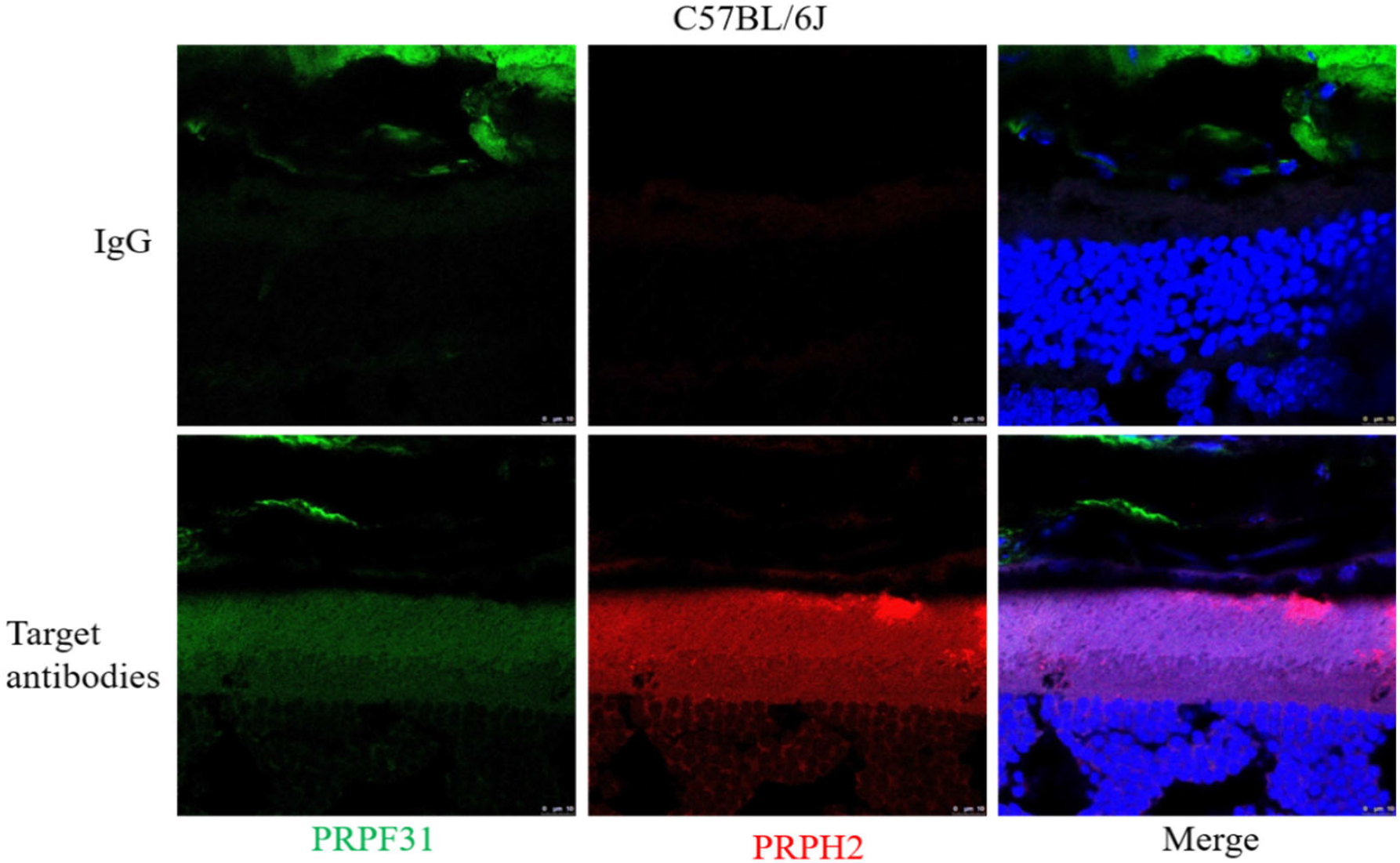
Immunohistochemistry showed that PRPF31 and PRPH2 partially co-localized in the photoreceptor of C57BL/6J mice. The green channel represents PRPF31, the red channel represents PRPH2, and the blue channel represents DAPI. The upper panel is IgG control without the primary antibodies, and the lower panel is with primary antibodies.

## Discussion

In this study, we showed partial co-localization of PRPF31 and PRPH2 in HEK293T cells, ARPE-19 cells and photoreceptors of C57 mice. Co-IP experiments showed an interaction between PRPF31 and PRPH2 in HEK293T cells and ARPE-19 cells.

The pathogenesis of retinitis pigmentosa remains not fully understood. Although there are 69 genes identified as the causative genes of retinitis pigmentosa and 280 genes for inherited retinal diseases on RetNet, more genes may be identified using more advanced genetic techniques. Furthermore, the function of these identified genes and their mutations should be investigated to understand the pathogenesis. PRPH2 is expressed in the rim regions of the flattened disks that contain rhodopsin of the outer segment of photoreceptors. Its function was reported as an adhesion molecule involved in the stabilization and compaction of outer segment disks. Pre-mRNA processing factor 31 homolog, PRPF31, is widely expressed in many tissues of the body. However, the mutation of PRPF31 only causes RP. It was reported that PRPF31 could bind to small nuclear ribonucleoprotein, snRNP, U5 and U4/U6 and regulate the processing of rhodopsin pre-mRNA. The detailed mechanism of the genes in RP remains to be further investigated.

The immunohistochemistry and immunocytochemistry showed that PRPF31 and PRPH2 co-localized in both HEK293T cells and ARPE-19 cells and suggested that they have a close spatial relationship. In the retina of C57BL/6 mice, PRPF31 and PRPH2 were expressed on the inner segment and outer segment of the photoreceptor. This suggests that both proteins participate in the balance of the outer segment. Further experiments using co-immunoprecipitation showed that PRPF31 could be pulled down by an anti-PRPH2 antibody and vice versa. When the PRPH2 protein was pulled down by the antibody, PRPF31 also precipitated. These results suggest that there is an interaction between PRPF31 and PRPH2.

There are some limitations in the current study. The cells used in this study are HEK293T cells and ARPE-19 cells. HEK293T is a human embryonic kidney cell line that differs from photoreceptors. Both PRPF31 and PRPH2 are also expressed on RPE cells. So, the ARPE-19 cells may be able to demonstrate the function of these genes in RPE. We did not use 661W cells, which is the cone photoreceptor cell line, but not rod [13]. Further studies are needed using the primary cell culture of the rod. Furthermore, the interaction of PRPF31 and PRPH2 was only demonstrated by co-immunoprecipitation but not by yeast two-hybrid (Y2H) screening systems. Because lots of proteins undergo varieties of post-translational modifications in mammal cells, which might lack in yeast. The failure of PRPF31and PRPH2 interaction assay in yeast might be reason of post-translational modifications difference. However, gene knockout or knockdown techniques are needed to further investigate the interaction in order to provide more strong evidence for their physiological interaction.

In conclusion, immunohistochemistry and immunoprecipitation suggest co-localization and interaction between PRPF31 and PRPH2. These findings may provide new insight into the pathogenesis of RP.

## Acknowledgment

This study was supported in part by Guangdong Nature Science Foundation (2018A0303130306).

